# Multi-layer Regulation of Rubisco in Response to Altered Carbon Status in *Synechococcus elongatus* PCC 7942

**DOI:** 10.1101/2021.10.11.463961

**Authors:** Amit K. Singh, María Santos-Merino, Jonathan K. Sakkos, Berkley J. Walker, Daniel C Ducat

## Abstract

Photosynthetic organisms possess a variety of mechanisms to achieve balance between absorbed light (source) and the capacity to metabolically utilize or dissipate this energy (sink). While regulatory processes that detect changes in metabolic status/balance are relatively well-studied in plants, analogous pathways remain poorly characterized in photosynthetic microbes. Herein, we explore systemic changes that result from alterations in carbon availability in the model cyanobacterium *Synechococcus elongatus* PCC 7942 by taking advantage of an engineered strain where influx/efflux of a central carbon metabolite, sucrose, can be regulated experimentally. We observe that induction of a high-flux sucrose export pathway leads to depletion of internal carbon storage pools (glycogen), and concurrent increases in photosynthetic parameters. Further, a proteome-wide analysis and fluorescence reporter-based analysis revealed that upregulated factors following the activation of the metabolic sink are strongly concentrated on ribulose-1,5-bisphosphate carboxylase-oxygenase (Rubisco) and axillary modules involved in Rubisco maturation. Carboxysome number and Rubisco activity also increase following engagement of sucrose secretion. Conversely, reversing the flux of sucrose by feeding exogenous sucrose heterologously results in increased glycogen pools, decreased Rubisco abundance, decreased photosystem II quantum efficiency, and carboxysome reorganization. Our data suggest that Rubisco activity and organization are key outputs connected to regulatory pathways involved in metabolic balancing in cyanobacteria.

**One sentence summary:** Rubisco activity and organization are key outputs involve in source-sink balancing in cyanobacteria.

## Introduction

Photosynthetic organisms require regulatory mechanisms to overcome dynamic fluctuation in solar illumination (*e*.*g*., diurnal cycle, sun-flecks), with an ultimate goal of aligning light energy inputs (“source”; *i*.*e*., absorbed photonic energy not dissipated by photoprotective mechanisms) with an equivalent capacity to utilize this energy using anabolic metabolism (“sinks”) (White et al., 2016; Walker et al., 2020). Adaptive responses that poise light harvesting antenna to a given light environment (Montgomery, 2014; Ruban and Belgio, 2014) and photoprotective processes that dissipate or redistribute excess light excitation (Allahverdiyeva et al., 2013; Roach and Krieger-Liszkay, 2014; Gerotto et al., 2016) have been relatively well-interrogated in cyanobacteria. Additionally, in plants, key metabolite pools (*e*.*g*., sucrose, trehalose-6-phosphate) are monitored as a proxy of source/sink energetic status and integrated into signalling networks that poise the expression of photosynthetic machinery, including photosystems, light harvesting antenna, and ribulose-1,5-bisphosphate carboxylase-oxygenase (Rubisco) (Mccormick et al., 2009; Adams et al., 2013; Lemoine et al., 2013; Sakr et al., 2018; Roth et al., 2019b; Santos-Merino et al., 2021). By contrast, cyanobacteria lack homologs for these signalling functions, making it uncertain how they sense the integrated metabolic demands required for cell homeostasis/growth, and coordinate upstream photosynthetic machinery to meet those energetic needs.

A key bottleneck between the light reactions of photosynthesis and downstream central carbon metabolism is Rubisco: a hexadecameric protein complex of large (RbcL) and small (RbcS) subunits that catalyze the carbon fixation step of the Calvin-Benson-Bassham (CBB) cycle and has notorious for its low catalytic activity and substrate specificity limitations (Spreitzer and Salvucci, 2002; Tcherkez et al., 2006). To overcome Rubisco’s enzymatic limitations, cyanobacteria depend upon carbon concentrating mechanisms (CCMs). The cyanobacterial CCM is distinguished by unique features including the carboxysome, a subcellular compartment that greatly enhances the local concentration of CO_2_ near Rubisco (Yeates et al., 2008; Borden and Savage, 2021). Structurally, the carboxysome is a protein microcompartment with an outer coat consisting of various of protein shell forms (hexamer, pentamer and trimer), while the inside of the compartment is packed with a liquid like matrix containing the protein CcmM, which acts to condense Rubisco into a paracrystalline-like array (Cameron et al., 2013; Wang et al., 2019). Plasma membrane-localized bicarbonate transporters actively pump inorganic carbon into the cytosol, which is thought to diffuse through selective carboxysome shell pores, whereupon it is converted into CO_2_ by encapsulated carbonic anhydrase. The end result is a concentration of CO_2_ around Rubisco up to ~1,000-fold higher than ambient levels (Badger and Price, 2003).

The cyanobacterial CCM is dynamically regulated in response to environmental changes (Raven and Beardall, 2014), a feature that appears to be important for cyanobacterial adaptation to a wide range of ecosystems. Environmental cues (light, CO_2_ and temperature) impact bicarbonate transporter gene expression, Rubisco content, and carboxysome composition/morphology (Logothetis et al., 2004; Mackenzie et al., 2004; Sun et al., 2016; Jahn et al., 2018; Rillema et al., 2020). The size and number of carboxysomes has been shown to be regulated in response to light quality and quantity (Rohnke et al., 2018; Sun et al., 2019). These changes in carboxysome structure are predicted to impact the relative capacity of the compartment to scavenge and uptake CO_2_ under different contexts (Mangan and Brenner, 2014). The emerging picture suggests that cyanobacteria regulate the carboxysome to modulate Rubisco activity and optimize cell growth and carbon fixation to environmental conditions. Yet, how CCM regulation is integrated with metabolism and/or changing metabolic demands (*e*.*g*., total metabolic flux/load) remains relatively unexplored.

In plants, signalling pathways act to control the activity of Rubisco in response to downstream metabolic status and to achieve source/sink balance. One well-conserved example involves Hexokinase family members that sense key metabolite pools of carbohydrates. For example, Arabidopsis Hexokinase 1 (HXK1) translocates into the nucleus following binding of glucose and forms a complex that suppresses expression of photosynthesis genes including the Rubisco small subunit, chlorophyll *a* (Chl *a*), and carbonic anhydrase (Cho et al., 2006). HXK1 has recently been shown to have conserved roles in the regulation of carbon balance in the microalga *Chromochloris zofingiensis*, and is implicated in a rapid change in the transcriptome of nearly a third of the total genome when cells are fed exogenous sugars, including many genes involved in photosynthesis, Chl *a* biogenesis and Rubisco maturation (Roth et al., 2019a).

Recent studies in cyanobacteria support the hypothesis that activation of heterologous metabolic pathways (*e*.*g*., engineered bioproduction circuits) can redistribute cellular resources in a manner that requires energetic re-balancing, including that of upstream photosynthetic processes. For example, we have previously shown that in the hours following the activation of a sucrose-secretion pathway via the overexpression of sucrose phosphate synthase (SPS) and sucrose permease (CscB) in *Synechococcus elongatus* PCC 7942 (*S. elongatus*) that there is a notable increase in the relative flux through the photosynthetic electron transport chain (PET), enhanced carbon dioxide fixation rates, and reduced acceptor-side limitations on the activity of PSI (Ducat et al., 2012; Abramson et al., 2016; Santos-Merino et al., 2021). We observe that the upregulation of photosynthetic flux is proportional to the amount of cellular resources that are redirected to the heterologous pathway (Abramson et al., 2016; Santos-Merino et al., 2021); *i*.*e*., when up to ~80% of photosynthetically fixed carbon is rerouted to the secreted sucrose bioproduct) More widely, a number of other cyanobacterial species and strains engineered to export other carbon metabolites have been shown to experience similar photosynthetic enhancements when heterologous metabolism is engaged, including isobuteraldehyde (Li et al., 2014), 2,3-butanediol (Oliver et al., 2013), and ethylene (Ungerer et al., 2012).

In the present study, we have used sucrose exporting cyanobacterial strains (*S. elongatus* overexpressing both CscB and SPS; hereafter “CscB/SPS”) well-studied for the photosynthetic changes induced by the expression of their heterologous carbon pathway (Ducat et al., 2012; Abramson et al., 2016; Santos-Merino et al., 2021), as a system that allows experimental control over cyanobacterial sink energy balance. We undertook a systems-level analysis of proteomic changes that accompany engagement of the sucrose ‘sink’, finding that the most significant hits were concentrated around Rubsico and Rubisco-associated factors. Further analysis using live cell imaging and biochemistry shows that carboxysome and Rubisco abundance is dynamically regulated in response to expression of this heterologous sink. Sucrose-feeding experiments, whereby exogenous also supports a model whereby carboxysome number and organization is linked to metabolic status.

## Results

### System level proteomic response to the sucrose export

We sought to gain deeper insight into the adaptive cellular response that results from engagement of a strong heterologous carbon sink. We first validated that the previously described CscB/SPS strain (Abramson et al., 2016; Abramson et al., 2018; Santos-Merino et al., 2021) was capable of sucrose export under our experimental setup (Fig. 1b) and exhibited the previously-described increase in photosynthetic performance upon activation of this heterologous sink via IPTG addition (Supplementary Fig. S1). We also monitored the internal glycogen content in CscB/SPS 24-48 hours following IPTG induction, observing a decline in glycogen stores by >75% on a per-cell basis (Fig. 1C): at later time points, glycogen content partially recovers relative to non-secreting controls (~30% decrease at 120h). In other contexts, such as diurnal cycles, glycogen content positively correlates with cellular carbon abundance (Diamond et al., 2015), suggesting the decrease in glycogen content linked to sucrose export may be a function of increased carbon flux towards the heterologous sucrose export pathway. However, it should be noted that the rate of sucrose efflux is approximately 2 orders of magnitude larger than could be accounted for mobilization of glycogen stores alone (Fig. 1B, C).

**Figure 1.**
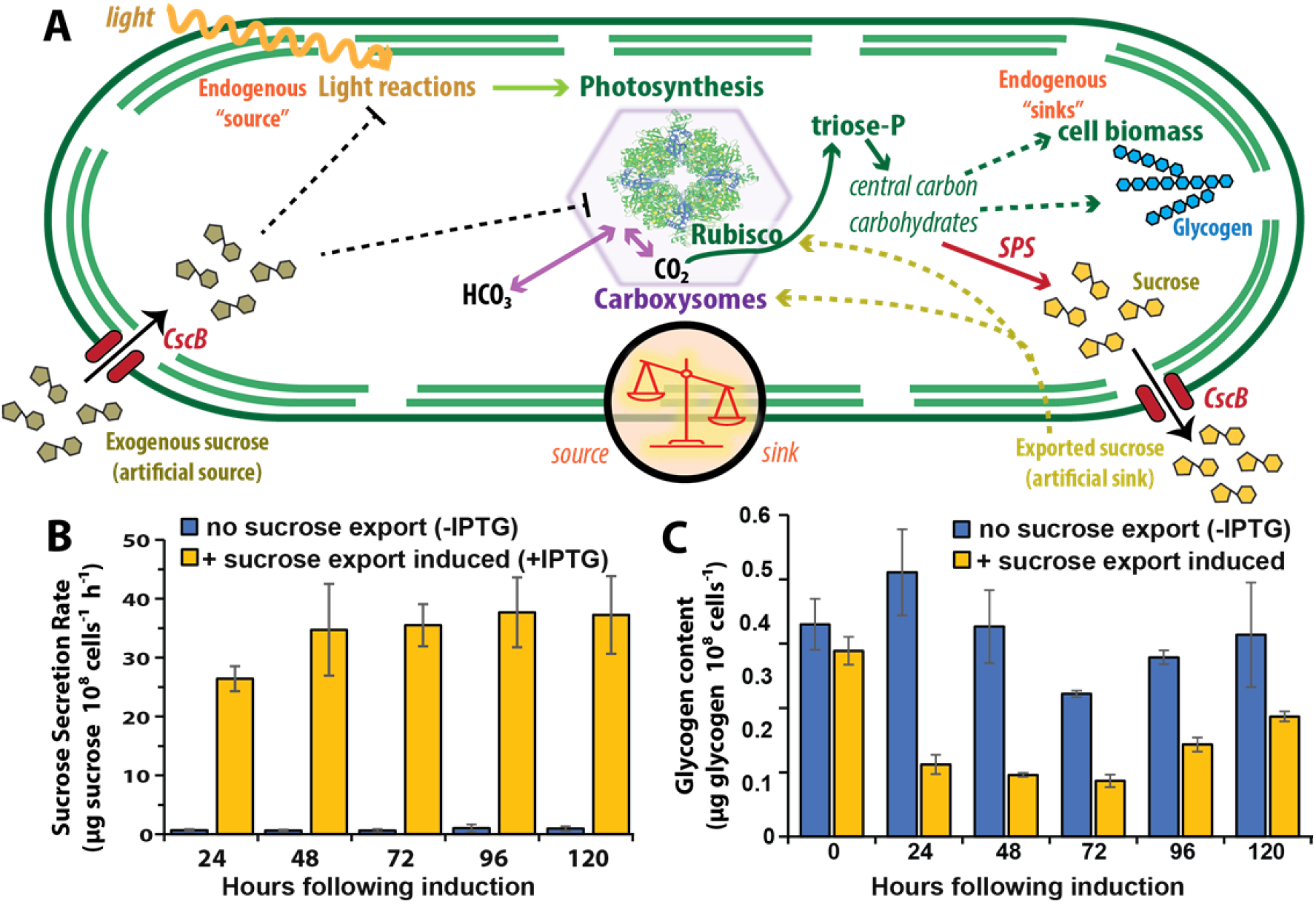
Connection of heterologous sucrose pathway with endogenous metabolism in *S. elongatus*. **A.** Schematic representation of cyanobacterial source/sink relationships. Endogenous metabolic sinks include metabolism leading to formation of glycogen, sucrose, and other cell biomass and depicted on the right. Cell inputs include light captured for photosynthesis as well as artificially supplied sucrose imported through heterologous transporters. Overexpressed genes for sucrose synthesis (sucrose phosphate synthase: SPS) and transport (sucrose permase: CscB) are shown in red. Quantification of exported sucrose (**B**) and internal glycogen stores (**C**) for strains induced to express SPS and CscB are shown in time series, with uninduced controls. Error bars represent the standard deviation of three independent biological replicates from the average in a representative time series.

To expand our analysis of the systemic changes following activation of a heterologous metabolic pathway, we chose an unbiased proteomic approach of CscB/SPS at time intervals 24, 48, 72 and 96h following induction of sucrose export. We unambiguously identified 913 proteins consistently detected across all sample conditions and timepoints (Fig. 2A), corresponding to a coverage of 34% of total proteins encoded in the genome of *S. elongatus* (913/2,657). Enrichment analysis using KEGG-assigned gene ontology terms allowed calculation of the percentage enrichment of identified proteins across 17 functional categories relative to the total gene products encoded in the genome (Fig. 2B). Across annotated functional categories, >50% proteins within each functional group were identified (Fig. 2B). A lower coverage of lower abundance and/or poorly characterized proteins was observed (240/913 proteins in our proteomics dataset).

**Figure 2.**
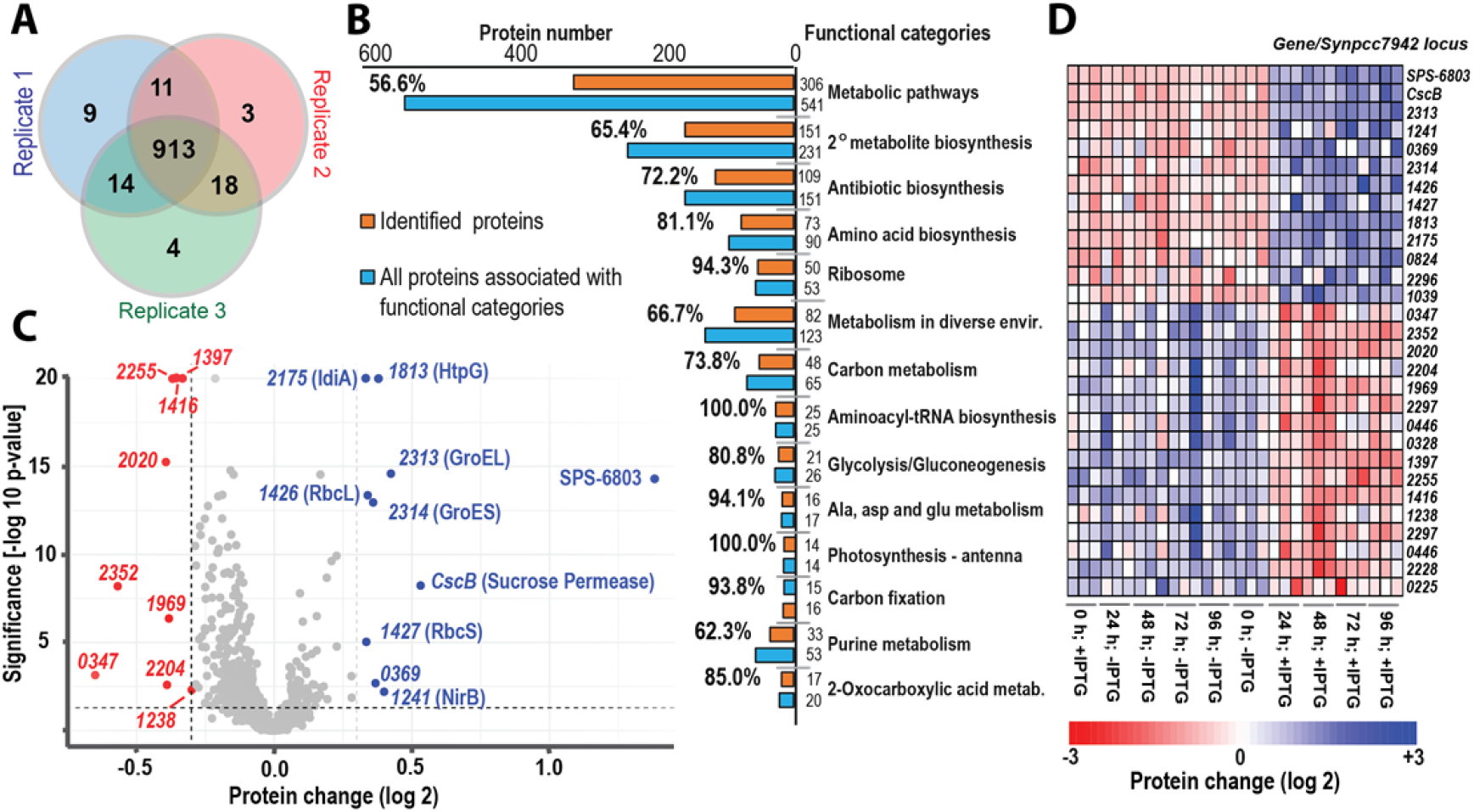
System level proteomic response to sucrose export. **A**. Venn diagram summarizing the number of unambiguously identified proteins inform untargeted proteomic analysis of three biological replicates. **B**. Representation of proteins from proteomic analysis within established annotated functional groups, with the number and percentage of identified proteins from the indicated categories relative to the total number of proteins with that designation in *S. elongatus* 7942, as assigned by KEGG pathway using STRING database resource. **C**. Volcano plot indicating changing protein abundance in induced strains integrated across time points 24, 48, 72 and 96 hours relative to controls. The non-axial vertical dashed-line shows ±0.3 log_2_fold protein change and non-axial horizontal dashed-line shows Mann-Whitney test p<0.05 with Benjamini-Hochberg correction cuttoffs. Differential protein analysis cut-offs were −1.3 > down-regulation and +1.3 < upregulation. Proteins represented by blue data points indicate significantly up-regulated proteins, while red points are downregulated. Proteins are identified with the abbreviated number ‘XXXX’ instead of the full genomic locus name (*i*.*e*., Synpcc7942_XXXX; Supplementary Table S1). **D**. Heatmap of significantly up- and downregulated proteins for each sample in the proteomic time-series.

Proteomic analysis identified a relatively small set of changes in protein abundance that were consistent across all time points following sucrose export (Fig. 2D). Importantly, among the most significantly upregulated proteins (10) identified in our proteomic analysis were CscB and SPS, the factors specifically induced in our engineered strains to trigger sucrose export (Supplementary Table S1). Among the most strongly upregulated endogenous proteins, the Rubisco enzyme subunits (RbcL and RbcS) were two of the protein which showed most statistically significant changes (Fig. 2C). Further, factors involved in the maturation of Rubisco were also over-represented among the most strongly upregulated proteins following sucrose export. This included chaperonins directly associated with the correct folding and assembly of Rubisco into higher order complexes, such as GroEL and its co-factor GroES which contribute to folding of RbcL and assemble of RbcL dimers (Hayer-Hartl et al., 2016). Another upregulated chaperone was HtpG (Synpcc7942_1813), a member of the heat-shock protein (Hsp90A) family which play roles in thermal or oxidative stress response in *Synechococcus* (Hossain and Nakamoto, 2003; Kobayashi et al., 2017). Three additional proteins that were identified as strongly upregulated were; IdiA, a factor associated with protection of photosystem II under various stress conditions (Yousef et al., 2003); Synpcc7942_0369, a conserved but poorly characterized putative oxidoreductase, and; NirB, a protein involved in nitrate assimilation and carbon/nitrogen balance (Ohashi et al., 2011) (Fig. 2C).

The seven proteins that were significantly downregulated following engagement of the sucrose export pathway were not as clearly concentrated around a common molecular function (Supplementary Table S1). Three of the downregulated targets were subunits related to ribosomal activities (Synpcc7942_2020, Synpcc7942_2352, and Synpcc7942_2204) (Hood et al., 2016). Two proteins in the AbrB–like family (Synpcc7942_1969 and Synpcc7942_2255) were downregulated, these have been characterized to act as transcription factors involved in carbon/nitrogen balancing in *Synechocystis* PCC 6803 (Lieman-Hurwitz et al., 2009; Yamauchi et al., 2011; Orf et al., 2016; Rachedi et al., 2020). To better visualize the overall changes in the proteome, we mapped identified proteins with conserved and well-established molecular functions onto a proteomap (Supplementary Fig. S2), which can provide a crude approximation of the relative protein abundance of each factor as a function of the summation of identified peptides for each protein (Liebermeister et al., 2014). In addition to the aforementioned increase in abundance for subunits of Rubisco following sucrose export, the proteomap highlights a subtle decrease in abundance across a number of proteins involved in ribosomal function and components of the photosynthetic electron transport chain (Supplementary Fig. S2).

### Rubisco is strongly upregulated and reorganized following sucrose export

To independently confirm proteomic analysis, we evaluated total Rubisco enzyme activity in cell extracts from sucrose exporting cells relative to controls. A significant increase in total Rubisco activity was observed at all time intervals when normalized to Chl *a* content, peaking at ~50% increased activity at 48h post-induction (Fig. 3A). Quantitative Western blots also indicated an increase in Rubisco levels following induction of the sucrose secretion pathway (Fig. 3B), this increases similar in magnitude to the enhanced activity of the *in vitro* enzyme assay. Increased Rubisco activity is consistent with prior reports that have shown that total carbon fixation rates and total biomass accumulation (*i*.*e*., cell biomass plus secreted carbon biomass) increase on a per cell basis when the sucrose secretion pathway is induced (Ducat et al., 2012; Abramson et al., 2016; Santos-Merino et al., 2021).

**Figure 3.**
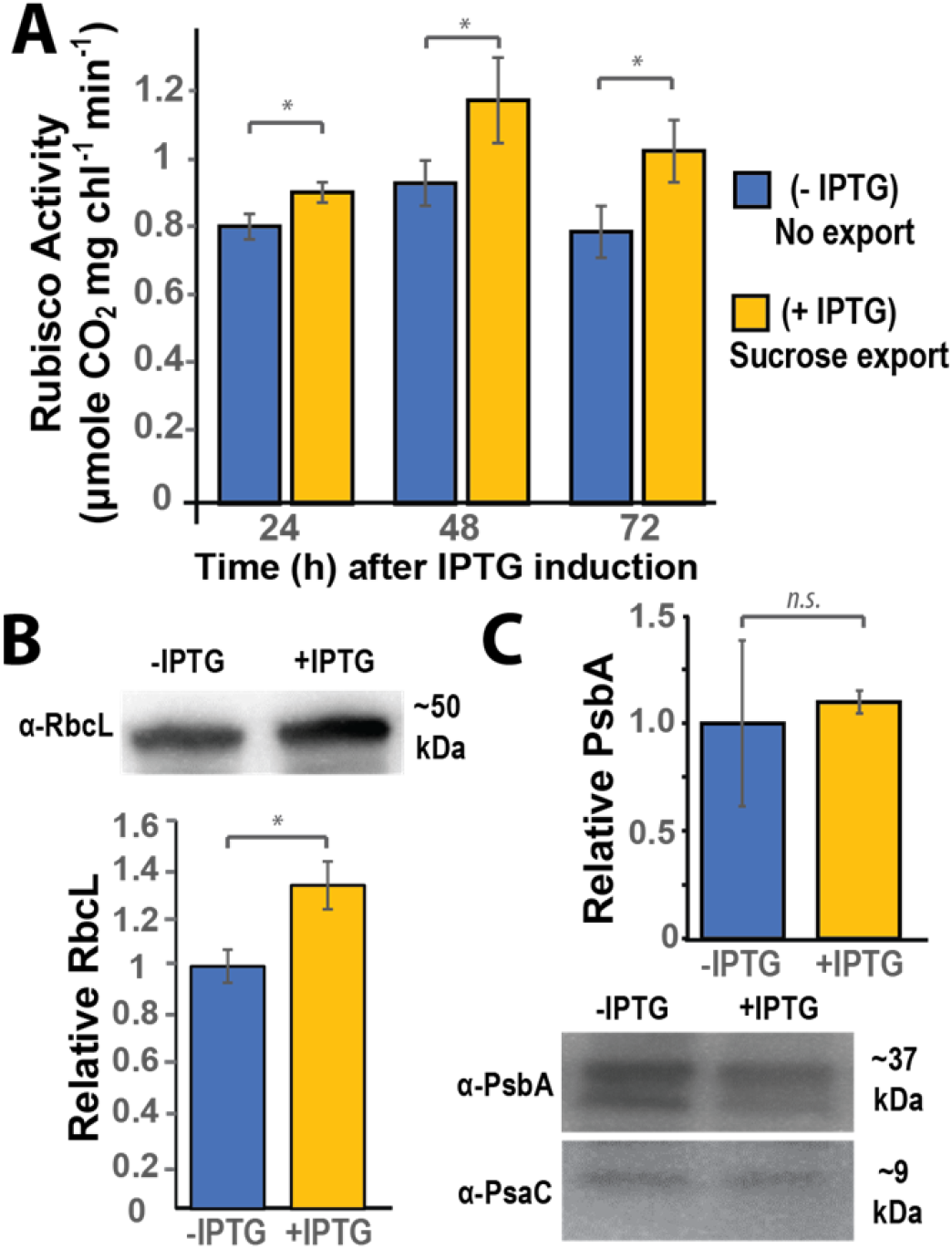
Rubisco is strongly upregulated and reorganized following sucrose export. **A**. Rubisco activity was measured from CscB/SPS lysates at 24h interval following induction of sucrose export (+IPTG) in comparison to uninduced controls (−IPTG). The activity measured for each sample was normalized to the Chl *a* content of the same sample. Western blots of (**B**) RbcL (**C**) PsbA, or PsaC levels were examined 72 hours post-induction (+IPTG) and normalized against uninduced controls (−IPTG). Error bars indicate the standard deviation of three independent biological replicates. Asterisk ‘*’ indicates statistical significance (p<0.05) against controls.

While we have previously reported enhancements in photosynthetic performance within the hours following induction of the heterologous sucrose sink (Supplementary Fig. S1 and (Ducat et al., 2012; Abramson et al., 2016; Santos-Merino et al., 2021), we do not find comparable evidence for significant alterations in the abundance of the light harvesting machinery. Consistent with our proteomics dataset (Fig. 2C and Supplementary Fig. S2), quantitative Western blotting for subunits of key components of the photosynthetic electron transport chain does not show substantial changes in representative subunits of PSI (PasC) and PSII (PsbA) (Fig. 3C).

### Effect of exogenous sucrose on photosynthetic activity and glycogen content

One possible interpretation of the physiological changes observed following induction of the sucrose export pathway is that they may be partially related (directly or indirectly) to the depletion of internal pools of carbon and/or energy equivalents. Pathways involved in carbon/energy sensing in plants were classically identified by manipulations that increased or decreased flux of carbon (*e*.*g*., sucrose) to specific tissues (Rolland et al., 2006; Lemoine et al., 2013). In order to determine if artificial increases in carbohydrate availability would impact similar cellular features, we examined the effect of supplying exogenous sucrose on cell physiology (Fig. 1A). We first analyzed the impact of sucrose feeding on Chl *a* and stored glycogen by varying the concentration of sucrose (0-200mM) with or without inducing expression of the sucrose transporter (CscB). Glycogen content increased (by 2-3 fold) in cells where external sucrose was supplied and CscB was induced to facilitate sucrose uptake in *S. elongatus* (Fig. 4A). In the absence of exogenous sucrose (*i*.*e*., 0mM sucrose), inducing expression of CscB did not change glycogen pools. A small, but significant increase in glycogen content was observed at higher exogenous sucrose concentrations (150-200 mM), possibly indicating alternative uptake pathways or indirect effects of the increased osmotic pressure (Page-Sharp et al., 1998; Suzuki et al., 2010).

**Figure 4.**
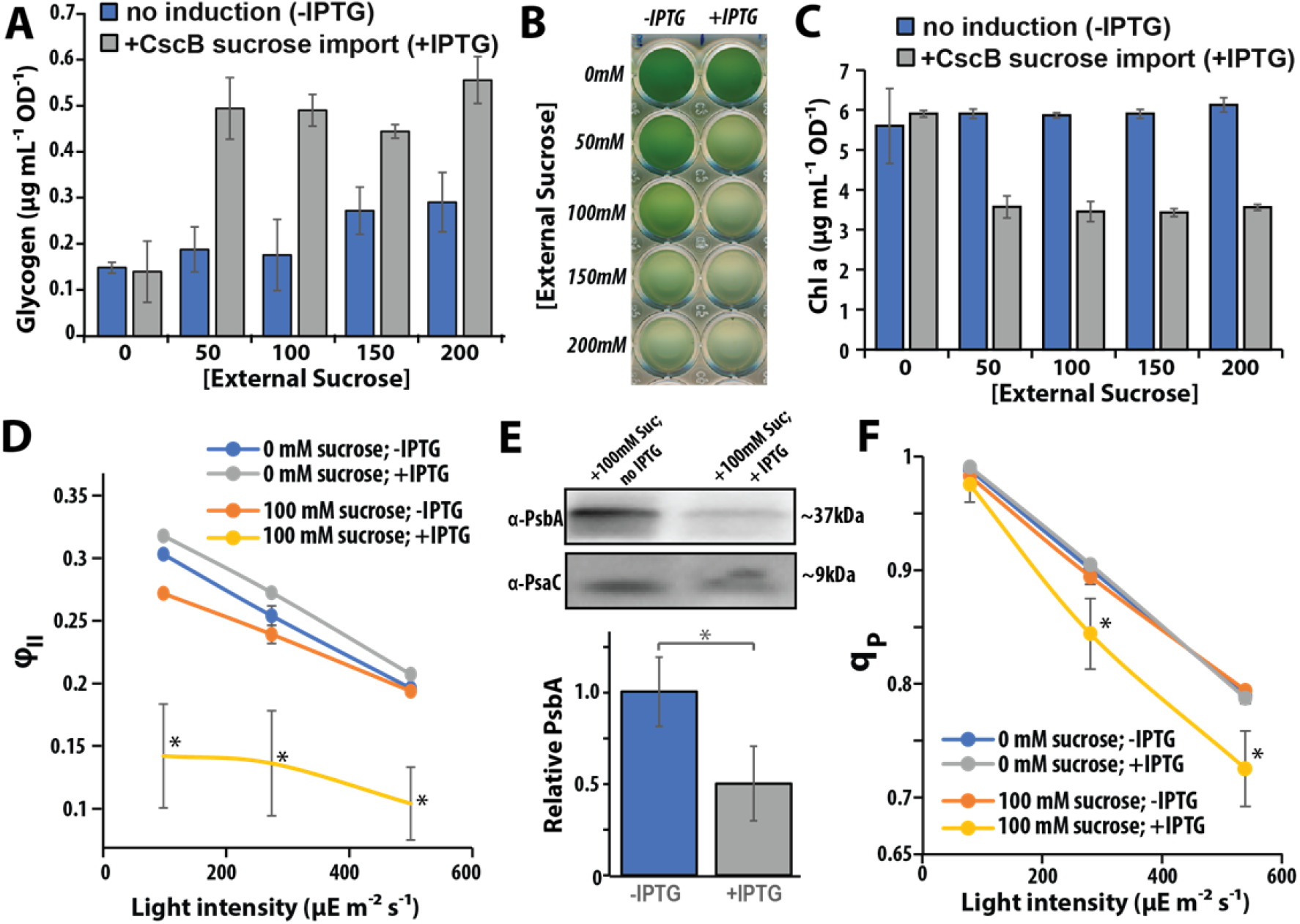
Effect of sucrose uptake on photosynthetic activity and glycogen content. External sucrose was supplemented in growth medium at indicated levels and CscB mutant lines were induced to allow uptake through sucrose permase expression. **A**. Effect of external sucrose uptake on glycogen content. Measurements of glycogen content were observed under induced (+IPTG) and uninduced (Control; −IPTG) at 24h time interval. **B**. Visual bleaching of CscB strains at 24h following sucrose uptake. **C**. Chl *a* content of cultures incubated with external sucrose at 24 h. Photosynthetic parameters such as PSII quantum yield (ϕ_II_) **(D)** and photochemical quenching (q_p_) **(F)** were analysed using Chl *a* fluorescence-based spectrometer at actinic light intensities of 100, 275 and 500 μE m^-2^ s^-1^. **E**. Western blot of PsbA, PsaC and relative signal density PsbA abundance were analyzed in induced and uninduced conditions supplemented with external sucrose. Error bars indicate standard deviation of ≥3 biological replicates and asterisk ‘*’ shows significant level with p<0.05.

Chl *a* content was strongly anti-correlated on a per-cell basis with uptake of exogenous carbohydrates, as treatment with 50-200mM sucrose caused chlorosis only in strains with induced CscB (Fig. 4B,C). As a rise in the total photosynthetic activity and quantum efficiency of PSII are among the most well-characterized aspects that follow secretion sucrose (Ducat et al., 2012; Abramson et al., 2016; Santos-Merino et al., 2021) and other carbon products (Oliver et al., 2013), we first evaluated the impact of sucrose feeding on measures of quantum yield of PSII (φ_II_) and PSII openness (q_P_). The maximum quantum yield of PSII was significantly reduced under conditions of sucrose uptake across all actinic light intensities (Fig. 4D). PSII openness, as estimated by q_P_, declined at high actinic light intensity (275 and 500 μmol photons m^-2^ s^-1^) in IPTG induced cultures supplemented with 100mM sucrose, possibly indicative of an over-reduced plastoquine (PQ) pool relative to controls (Fig. 4F).

In tandem with the decrease in metrics of photosynthetic performance, we observed downregulation of some other components involved in the PET and associated light harvesting machinery. Quantitative western blot analysis indicated an ~2-fold decrease in the abundance of a core PSII subunit, PsbA, in the presence of 100mM when CscB was expressed (Fig. 4E). A central PSI subunit, PsaC, also appeared to be reduced in abundance (Fig. 4E; bottom), albeit to a lesser extent than PsbA. Phycobilisomes a major component of light harvesting antennae composed of two phycobiliproteins, allophycocyanin and phycocyanin, were also reduced by ~72% and 70% in sucrose fed cells, respectively (Supplementary Fig. S3).

### Carboxysomes are reorganized in response to sucrose export and uptake

In cyanobacteria, the bulk of Rubisco is housed within the lumen of the carboxysome. Since we observed changes in Rubisco activity/abundance following sucrose export and import, we examined if the cyanobacterial carboxysome was also altered in response to these interventions. To visualize changes in carboxysome organization, we expressed an exogenous copy of the small subunit of Rubisco fused to the fluorescent reporter mNeonGreen (RbcS-mNG) under the control of the native *P*_*rbcLS*_ promoter. Similar constructs have been employed by our group and others to examine carboxysome dynamics *in vivo* (Savage et al., 2010; Cameron et al., 2013; Chen et al., 2013; Hill et al., 2020). As expected, the RbcS-mNG reporter was concentrated to carboxysomal foci that were arranged along the central axis along the length of the cell when expressed in the mutant background of our strains containing the exogenous *cscB* and/or *sps* genes (Fig. 5A and Supplementary Fig. S4). Carboxysomes were most frequently arranged in a linear or hexagonal packing that maximizes the distance between neighboring microcompartments (Maccready et al., 2018).

**Figure 5.**
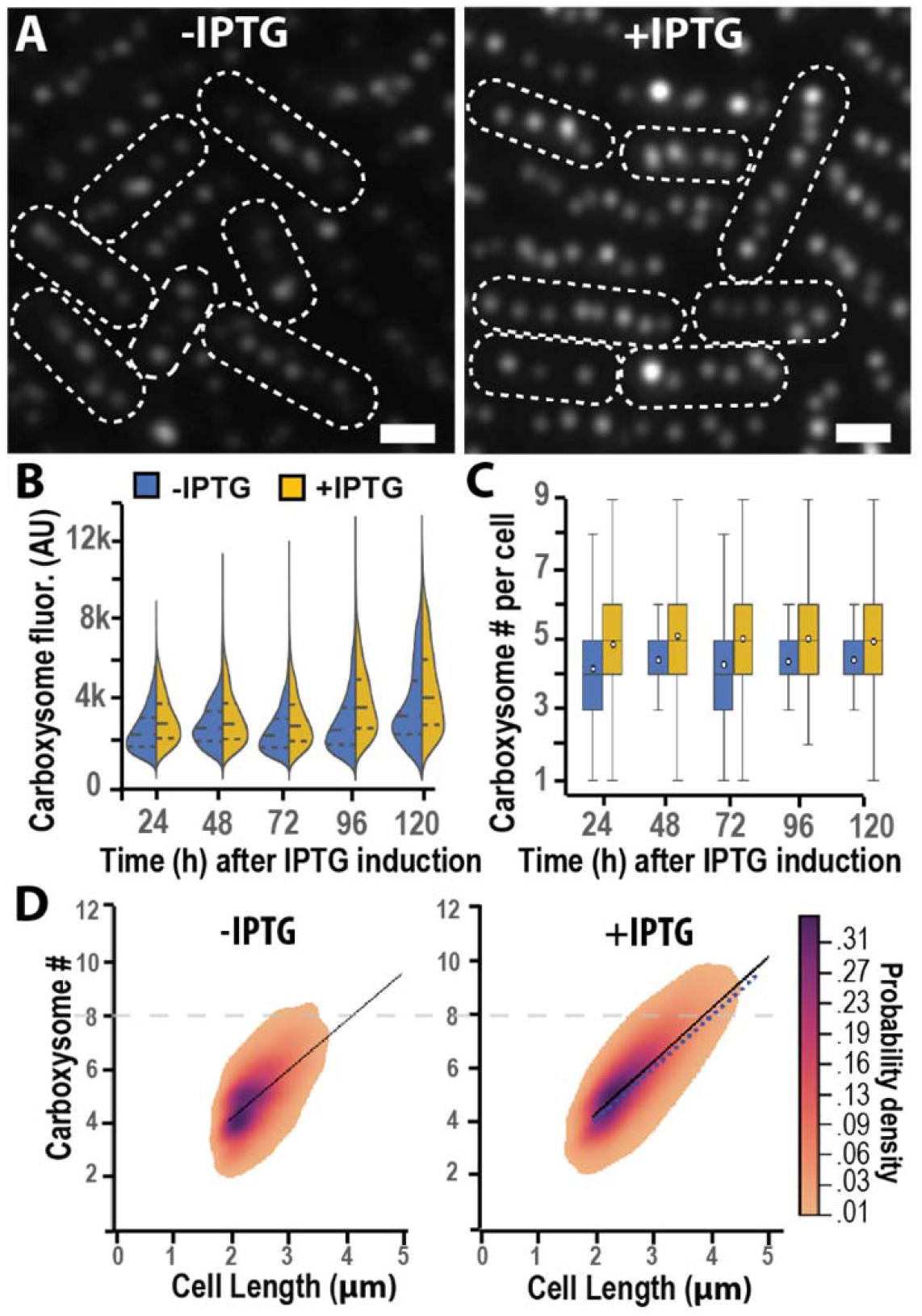
Carboxysomes are reorganized in response to sucrose export. *S. elongatus* lines bearing a RbcS-mNG reporter were used to visualize carboxysome organization following induction of sucrose export pathway. **A**. Representative images of CscB/SPS strains 72h after sucrose export induction (+IPTG) or control treatment (−IPTG). Violin plots represent the distribution of carboxysome puncta fluorescence intensity (B) and number per cell (C) in induced (+IPTG) cells compared to uninduced controls (−IPTG). (**D**) Density plot of carboxysome number as a function of the containing cell length in both uninduced (−IPTG) and induced (+IPTG) conditions. Probability density is indicated by color, while a linear best-fit trendline is displayed. For visual comparison, a blue dotted trendline. Each mutant strains have n>4,000 cells. Scale bars = 1 µm.

When sucrose export was induced through the heterologous expression of CscB and SPS, we observed changes in carboxysome number and in the intensity of foci (Fig. 5B, C, Supplementary Fig. S4). The intensity of RbcS-mNG puncta was noticeably brighter within 24 hours of induction of sucrose export, and this difference was maintained for multiple days relative to uninduced controls (Fig. 5B,C). The number of carboxysomes contained within each cell was also increased in the hours following induction of sucrose export (Fig. 5B,C). Since a slight cell elongation and narrowing of cell width was also evident in sucrose-secreting cells (Fig. 5A, Supplemental Fig. S5), we quantified carboxysome density, observing a slight, but significant increase in carboxysome number relative to cell length in sucrose-exporting cells (Fig. 5D). The ratio of cytosolic RbcS-mNG to carboxysome RbcS-mNG remained constant (Supplementary Fig. S6). Taken together, this is in agreement with our prior evidence for increased Rubisco content in sucrose-secreting cells (Figs. 2, 3), and suggests that the additional Rubsico remains packaged within carboxysomes, resulting in increased carboxysome number and/or quantity of Rubsico per carboxysome.

When strains containing both the *rbcS-mNG* and *cscB* constructs were fed with exogenous sucrose, we observed changes in carboxysome organization that were tied to sucrose uptake (Fig. 6), as well as subtle changes in cell width (Supplementary Fig. S7). The number and density of carboxysomes declined in sucrose-importing strains (100mM Sucrose +IPTG; Fig. 6A and Supplementary Figs. S8, S9) and there was a change in carboxysome organization (Fig. 6A). We observed increased clustering of carboxysome puncta in many cells with the capacity to import exogenous sucrose (Fig. 6A (+IPTG); red arrowheads, Supplementary Fig. S8). We also observed an increase in the heterogeneity of puncta size/intensity within each cell, some of this may be attributable to clustering of carboxysomes, but in other cases puncta that appeared to be single carboxyxomes at the resolution limits of light microscopy also exhibited notably brighter RbcS-mNG fluorescence relative to puncta within the same cell (Fig. 6A,C; blue arrowheads). Direct assessment of Rubisco protein abundance and activity levels under sucrose feeding conditions in mutant background a strain containing CscB (but lacking the RbcS-mNG reporter) indicated a ~25% reduction in Rubisco content on a total protein basis (Fig. 6E) and a ~60% decline in Rubisco activity on a per-cell basis (Fig. 6B).

**Figure 6.**
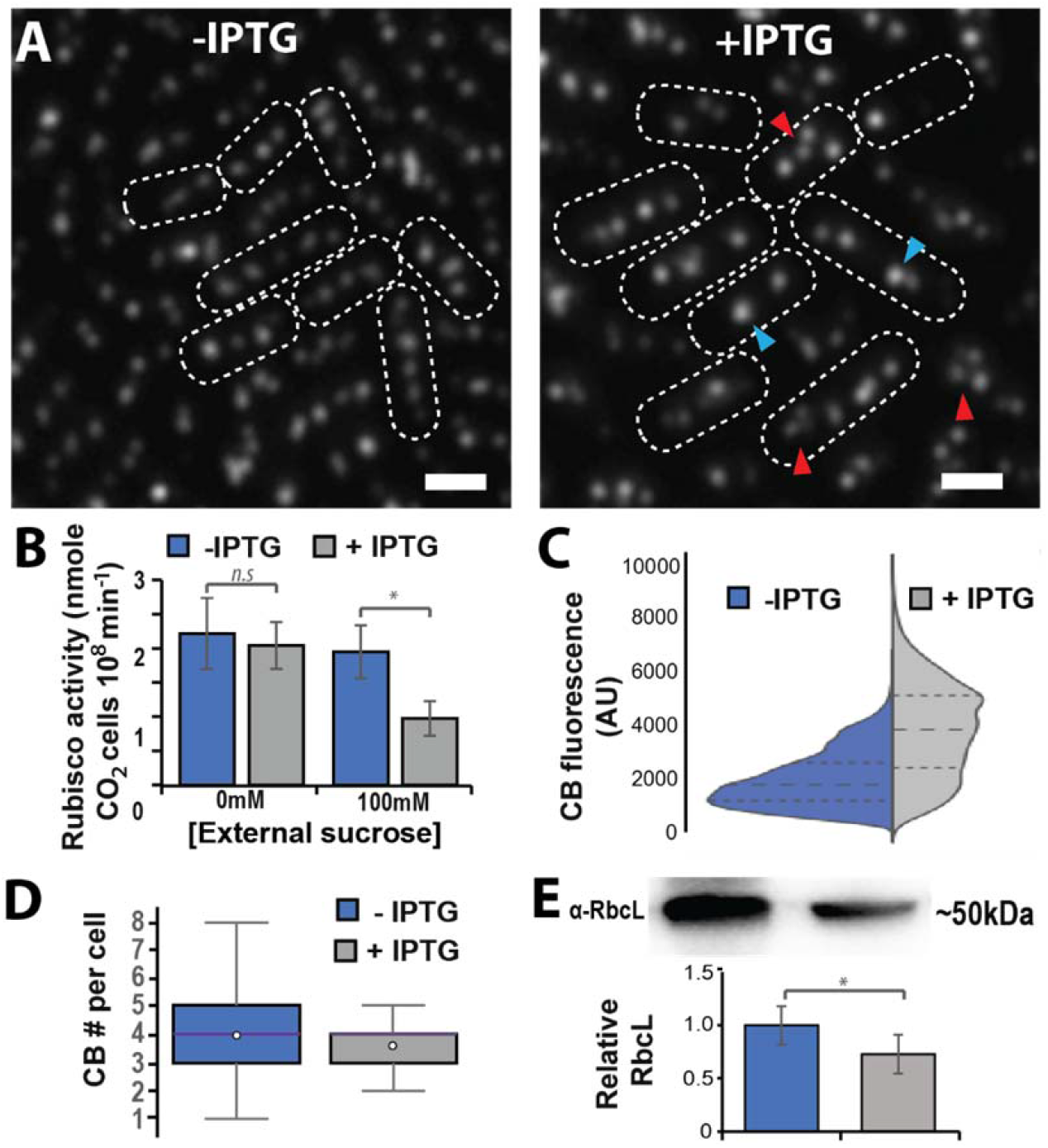
Carboxysomes are reorganized in response to sucrose import. **A**. Carboxysome reporter (RbcS-mNG) expressed in CscB line was visualized following 24h of incubation in 100mM external sucrose with the sucrose transporter induced (+IPTG) or without (−IPTG). **B**. Rubisco activity in in strains as above was measured after 24h of incubation the indicted external sucrose level. The activity measured for each sample was measured as a function of cell number. Violin and box plots depicting the difference in the distribution of carboxysome puncta intensity (**C**) and carboxysome number (**D**) for sucrose importing (+IPTG) strains relative to uninduced. Western blot analysis and relative signal density of rubisco large subunit (RbcL) abundance in induced and uninduced conditions **E**. Each bar represents the mean of three independent biological replicates (±SD). Asterisk ‘*’ shows significant level with p<0.05. Each mutant strains have n>4,000 cells. AU, Arbitrary units. Scale bars = 1 µm.

## Discussion

Our results suggest that cyanobacteria exhibit numerous changes in photosynthetic components following an engagement of a heterologous sucrose export (sink) pathway, and that these changes are concentrated around Rubisco. We observe changes in the abundance, activity, and organization of Rubisco within carboxysomes after inducing sucrose export, and that many of these effects are reversed when exogenous sucrose is supplied to induce a state of carbon overabundance. Taken together, our evidence suggests that the cyanobacterial CCM – Rubisco and carboxysome organization especially - may respond to internal metabolic signals and source/sink dynamics.

### Proteomic changes enhance carbon fixation capacity following engagement of a heterologous carbon sink

We found that a relatively small subset of proteins is significantly altered in response to the activation of our engineered sucrose export pathway. Aside from the expected strong increase in the proteins required for sucrose production (CscB and SPS), 4 of the remaining 8 significantly upregulated proteins were subunits of Rubisco (RbcL, RbcS) or molecular chaperones with established functions in Rubisco’s maturation process (GroES, GroEL; Fig. 2C). We validated the proteomic results in Rubisco abundance by Western blot (Fig. 3B) and Rubisco activity assays (Fig. 3A), indicating that adjusted Rubisco levels are a primary target of regulation following engagement of the heterologous sink. This result suggests that the increase in total CO_2_ fixation rates that has been reported to follow sucrose export in previous publications (Ducat et al., 2012; Santos-Merino et al., 2021) in part stems from increased Rubisco catalytic activity, and not solely from the observed increases in the activity and efficiency of light reactions of photosynthesis (Abramson et al., 2016; Santos-Merino et al., 2021). Long-term changes in the abundance of other components of photosynthetic metabolism (*e*.*g*., photosystems, light harvesting machinery, subunits of the PET machinery) following sucrose export are relatively subtle or insignificant by both proteomics approaches and targeted assays (Figs. 2, 3).

We also report alterations in the organization of carboxysomes following sucrose export, as observed by imaging of live cells bearing a fluorescent Rubisco reporter (Fig. 5). It is intriguing that despite the observed increase in carboxysome number/density in sucrose-secreting cells (Fig. 5C,D), we did not detect significant upregulation of other carboxysome components in our proteomic analysis (Figure 2C). One possible explanation is that the average size of carboxysomes may be increased in response to sucrose export and that changes in carboxysome surface area (and associated shell/structural proteins) are relatively minor in comparison to the expanded lumenal volume. Increased carboxysome size would be consistent with the increased fluorescence intensity we see of carboxysome foci in sucrose-secreting cells (RbcS-mNG reporter; Fig. 5B, C). However, the resolution limits of light microscopy preclude the elimination of alternative interpretations, including: i) low sensitivity of our proteomics approach to shell proteins and other carboxysome structural components (Long et al., 2005; Faulkner et al., 2017); ii) carboxysome components may be in excess supply the cytosol of *S. elongatus* under our growth conditions, or; iii) the RbcS-mNG reporter may not behave identically to native RbcS with regard to luminal packaging.

One hypothesis regarding the physiological changes we observe following engagement of the sucrose secretion pathway is that they are representative of a regulatory response to the altered energy/carbon balance that occurs when a substantial proportion of cellular resources are redirected towards non-native processes (Santos-Merino et al., 2019). We and others reported a significant decrease in glycogen content in sucrose-secreting cyanobacteria, possibly attributable to the redirection of carbon pools (*e*.*g*., glucose, fructose) away from endogenous sinks and to the engineered pathway (Fig. 1C; (Qiao et al., 2018; Lin et al., 2020). Indirect evidence for this interpretation has also been provided in the form of studies showing that glycogen synthesis is in competition with bioproduct synthesis: reducing the capacity to store cellular carbon as glycogen can improve bioproduction from a number of engineered pathway (Ducat et al., 2012; Davies et al., 2014; Wang et al., 2020) While heterologous metabolism can mitigate impairments in growth and photosynthesis observed in cyanobacterial strains with restricted flux towards endogenous sinks (Li et al., 2014; Abramson et al., 2016; Xiong et al., 2017; Cano et al., 2018; Díaz-Troya et al., 2020; Santos-Merino et al., 2021).

### Rubisco activity and cellular organization is impacted following carbohydrate feeding experiments

If redirection of internal carbon pools away from endogenous metabolism and towards secreted bioproducts results in changes sensed by regulatory machinery in cyanobacteria, it would be expected that interventions that artificially increase internal carbon resources might also result in phenotypic changes in these cellular features. Although *S. elongatus* is regarded as a strict photoautotroph, it can grow photo-mixotrophically when heterologous transporters are expressed (McEwen et al., 2016), a property we used to artificially increase intracellular carbon availability by importing sucrose through the heterologous transporter, CscB. Increased glycogen content and reductions in φ_II_ and q_P_ (Fig. 4A,D,F) are consistent with increased cellular carbon resources and an over-reduced PET. Sucrose feeding experiments also result in a rapid downregulation of many components of the photosynthetic machinery, including light harvesting antennae, photosystems, and Rubisco (Figs. 4 and 6). Carboxysome number is decreased, and their spatial positioning is disrupted in sucrose-feeding experiments (compare Figs. 5 and 6). One possibility is that metabolic changes that accompany the influx of exogenous carbon impact (directly or indirectly) the activity of Maintenance of carboxysome distribution AB proteins (McdA/McdB) involved in microcompartment positioning along the cyanobacterial nucleoid (Maccready et al., 2018). An alternative speculative hypothesis is that exogenous carbohydrate uptake leads to changes in the integrity and/or dynamic association of carboxysome shell proteins on the bacterial microcompartment, as we have recently observed similar phenotypes when we visualize carboxysomes in real-time following destabilization of components of carboxysome shell proteins (specifically, CcmO or CcmL)(Sakkos et al., 2021). Distinguishing between these possibilities will require further research with more directed approaches to interrogate carboxysome dynamics and shell integrity. Regardless, our observations raise the possibility that carboxysome properties may be tied to internal metabolic states as well as external environmental conditions (*e*.*g*., light, CO_2_, temperature) previously reported (Woodger et al., 2003; Sun et al., 2016; Rohnke et al., 2018; Sun et al., 2019; Rillema et al., 2020).

### Possible implications for source/sink regulatory machinery in cyanobacteria

Rubisco is one of the primary regulatory targets for source/sink regulatory systems in plants (Nielsen et al., 1998; Cho et al., 2006; Granot et al., 2013; Koper et al., 2021). In addition to the previously mentioned mechanistic connection between HXK1 and Rubisco expression, multiple studies in plants have correlated intracellular carbohydrate availability with the expression of Rubisco and other components of the photosynthetic machinery. For example, interventions that decrease sink capacity relative to source energy (*e*.*g*., exogenous feeding of sugars) lead to decreased Rubisco abundance across many crop plants (Moore et al., 1998; Nielsen et al., 1998; Kasai, 2008; Lobo et al., 2015). Our results suggest that this relationship between Rubisco abundance and source/sink dynamics is maintained in cyanobacteria, although the specific mechanisms for monitoring energetic balance that have been elucidated in plants (*e*.*g*., HXK1, SnRK, TOR) do not appear to be conserved. This apparent functional conservation may result from the relatively high burden rubisco synthesis places on photosynthetic organisms (*i*.*e*., energetically and in nitrogen requirements). Stated differently, minimizing the cellular burden of rubisco synthesis in a given environment be similarly important to organismal fitness as is acquiring sufficient carbon fixation capacity to meet metabolic demands.

A deeper understanding of the mechanisms that regulate source/sink balance in cyanobacteria is likely to have biotechnological implications given the potential for cyanobacteria as a “carbon neutral” production chassis to combat anthropogenic climate change (Sabine et al., 2004; Zhang et al., 2017; DeLisi et al., 2020). Future research is required to determine mechanisms of cyanobacterial source/sink sensing so that they can be leveraged to maximize CO_2_ fixation and photosynthetic efficiency in cyanobacteria.

## Methods

### Growth medium and culture condition

Cultures of *S. elongatus* mutant strains were grown in BG-11 (Sigma-Aldrich) medium buffered with 1g/L HEPES (pH 8.3 with potassium hydroxide). For routine cultivation of cultures, an Infors-Multitron with 250 μmol photons m^-2^ s^-1^ compact fluorescent (GRO-LUX^®^) lighting supplemented with 2% CO_2_ was used at 30°C with orbital shaking at 130 rpm. Cultures were maintained with a daily back-dilution to OD_750_ ~0.3 unless otherwise noted. The sucrose exporting (CscB/SPS) and importing (CscB) mutants were used as previously describe by (Abramson et al., 2016). Strains bearing heterologous genes under P_trc_ promoters (*i*.*e*., *cscB* and/or *sps*) were induced with 1mM IPTG as indicated. The carboxysome fluorescence reporter Rubisco small subunit (RbcS-mNG) expression was driven by promoter (P_rbcLS_) and genomically inserted into Neutral Site (NS1) (Clerico et al., 2007). All mutant selection were carried out with BG11 plates with appropriate antibiotic supplementation; Spectinomycin (100μg/mL) and Chloramphenicol (25 μg/mL). Axenic liquid cultures were maintained through supplementation of the same antibiotics, although antibiotics were removed at least 3 days prior to the experiments described.

### Total Protein extraction and LC-MS/MS analysis

For protein extraction, 50 mL of culture was centrifuged (6,000 *×g*, 15 min, 4°C), supernatants were discarded, and pellets were transferred to 50 mL tubes. All steps of protein extraction were performed at 4°C. The pellets were resuspended in 10 mL of a protein extraction buffer (50 mM Tris-HCl, pH 7.6, 10 mM MgCl_2_, 0.1% TritonX-100, 1X of Halt protease inhibitor cocktail, Thermo, USA). Cell disruption was performed by French press (AMINCO^®^) at 1100 psi. After homogenization, the samples were centrifuged (17,000 *xg*, 10 min, 4°C) using round bottom tubes to remove cell debris. The supernatant was transferred to a 50mL conical tube and 4 volumes 100% acetone was added, whereupon samples were stored overnight at −20°C for complete protein precipitation. The samples were re-suspended in resuspension buffer (10mM Tris-HCl, 5% SDS, 1% β-mercaptoethanol, pH~ 8.0) and further used for proteomics measurements using LC-MS/MS (for details see Supplementary Method S1).

Raw LCMS/MS data was further processed for protein identification and differential expression analysis through Scaffold software (version 4.11.1, Proteome Software Inc., Portland, OR). For protein identification, 1% false discovery rate (FDR) and minimum 2 unique peptides were specified as cut-offs to filter data for protein identification and analysis. A significance level of *p*<0.05 (Mann-Whitney Test) and Benjamin-Hochberg correction were applied in Scaffold for differential analysis. Enrichment analysis was performed with KEGG (Kyoto Encyclopedia of Genes and Genomes) pathways assigned functional categories using STRING v11 data-base (https://string-db.org). Proteomaps were developed through modification and customization of the online tool (https://www.proteomaps.net/) following user documentation and literature available (Liebermeister et al., 2014).

### Fluorescence microscopy and image analysis

For microscopy images, 1mL cells were pelleted down by centrifuge at 5,000g for 5 min and pellet was resuspended in 100μL of BG-11. A 3μL aliquot was transferred to a 3% agarose pad. The cells were allowed to equilibrate for ~5min before the pad was placed onto a #1.5 glass coverslip for imaging. Fluorescence Images were taken by a Zeiss Axio Observer D1 microscope (100×1.3NA) with an Axiocam 503 (mono-chrome) camera. Images were further processed for pixel-based data analysis using MicrobeJ (v5.12d) an image analysis plugin for ImageJ (Abràmoff et al., 2005; Schindelin et al., 2012). MicrobeJ, as previously reported and described Ducret et al., 2016), was used to measure carboxysome foci fluorescence and number based on reporter mNG fluorescence (Ducret et al., 2016). The main attributes for defining cell are as follows (Fit shaped; rod shaped; Length: 1.5-10; Width:0.3-1.5; Area: 1-max) and smoothed maxima foci determination by (Tolerance: 100; z-score:10; Intensity: 0-max). This automated image analysis assisted in removing experimenter bias relative to manual image analysis, however, not all carboxysomes were identified in some instances (*e*.*g*., due to low puncta fluorescence or focal plane artifacts).

### Rubisco activity assay

Rubisco activity was assayed *in vitro* spectrophotometrically by following the coupled conversion of NADH to NAD^+^ (Ruuska et al., 1999). For protein extraction, 10ml of culture was centrifuged (6000 *×g*, 15 min, 4°C) and supernatant was discarded. The pellets were resuspended in 500µL of a protein extraction buffer (50mM EPPS, pH 8.1, 1mM EDTA, 10mM DTT, 0.1%TritonX-100 and 1X of Halt protease inhibitor cocktail, Thermo, USA) and transferred to 1.5mL tubes. The cell disruption was performed by Sonicator (Fisher scientific) using 20cycle (30s On and 10s Off) and amplitude 45% at 4°C. After protein extraction, the solution was centrifuged (6200 *xg*, 10 min, 4°C) to remove cell debris. To fully activate rubisco, cell-free extracts were incubated at room temperature for 20 minutes in the presence of 15 mM NaHCO_3_ and 15 mM MgCl_2_. Following activation, 40uL of the lysate was mixed in a cuvette with 960μL of an assay buffer containing 100mM HEPES, pH 8.1, 20mM MgCl_2_, 1mM EDTA, 1mM ATP, 20 mM NaHCO_3_, 0.2 mM NADH, 30 mM ribulose-1,5-bisphosphate and a coupling enzyme cocktail containing 20U glyceraldehyde-3-phosphate dehydrogenase, 22.5 U 3-phosphoglyceric phosphokinase, 12.5 U creatine phosphokinase, 250U CA and 56U Triose-phosphate isomerase. The reaction was initiated by adding sample, and the rate of NADH oxidation was monitored at 340nm for 10 minutes using UV spectrometer (Agilent). Activity was calculated from the molecular extinction coefficient of NADH. To avoid potential rubisco inhibitors often found in commercial preparations, RuBP was synthesized and purified in-house and confirmed to have minimal “fall-over” kinetics on purified rubisco samples (Kane et al., 1994; Kane et al., 1998).

### Biochemical assay for sucrose and glycogen content

2 mL cultures were pelleted in a centrifuge at 6,200 *xg*. Pellets were processed for glycogen assay and the supernatant was sampled for sucrose assays. The glycogen assay was performed following the protocol Nakajima et al (2017)(Nakajima et al., 2017) with modifications. Briefly, pellets were resuspended in 200µL of 30% KOH and incubated in a 95°C in water bath for 2h. After incubation, 600uL absolute ethanol was added and further incubated at –20°C overnight. The next day, the suspension was centrifuged, and pellet was dried in a oven. Dried pellets were resuspended in ddH_2_O and analysed with a commercially available Glycogen assay kit (EnzyChrom™, USA). Sucrose quantification from culture supernatants was determined using the Sucrose/ d-Glucose Assay Kit (Megazyme: K-SUCGL) following the manufacturer’s instructions.

### Pigment analysis

Photosynthetic pigments were solubilized from cell pellets by incubation in 100% methanol for 30 min at 4°C. Chlorophyll *a* was measured spectrophotometrically following the protocols of Porra et al. (Porra et al., 1989). Briefly, the pigment suspension was centrifuged, and supernatant was used for absorption at 665nm. Chl *a* content was determined using an UV/Visible Spectrophotometer (Genesis 20, Thermo, USA). Phycobiliprotens were measured following the protocol of Zavrel et al. (Zavřel et al., 2018).

#### Fluorescence measurements of photosynthetic parameters

Fluorescence of photosynthetic parameters were measured on a customized fluorimeter/spectrophotometer (Hall et al., 2013) modified for liquid samples. A cuvette with sample was illuminated with a pulsed measuring beam [*λ* = 590 nm peak emission light-emitting diode (LED), Luxeon Z Color Line] and then illuminated at three different intensities of photosynthetically active radiation (PAR), 100, 275 and 500 μmol photons m^−2^ s^−1^ (*λ* = 460 nm peak emission, Luxeon Rebel Royal-Blue LED). To acclimate the sample and minimize the impact of successive saturating pulses. the cuvette was illuminated at the relevant actinic light for 3 minutes and 2 minutes, respectively, before the first saturating pulse, and between each pulse. Cyanobacterial samples containing 2.5 μg mL^−1^ Chl *a* were pelleted, then resuspended in fresh medium sparged with 2% CO_2_ in air and dark-adapted for 3 min before fluorescence measurments. The relative yields of Chl *a* fluorescence were measured under steady-state illumination (Fs), during a 1.5 s saturating pulses of actinic light (~5,000 μmol photons m^−2^ s^−1^) (F’m) and after exposure to ~2 s of darkness with far-red illumination (F’_o_). Chl *a* fluorescence was used to calculate the quantum yield of PSII (Φ_II_), the coefficient of photochemical quenching (q_p_) using equations (1) and (2), respectively (Campbell et al., 1998; Maxwell and Johnson, 2000).

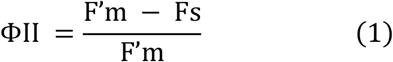

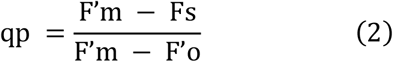

Where, F’m = the value of maximal fluorescence in the light-adapted state,Fs = the steady-state fluorescence in the light-adapted state, and F’o = the minimal fluorescence in the light-adapted state

### Western blot analysis

For Western blotting, cells were lysed in extraction buffer (50 mM Tris-HCl, pH 7.6, 10 mM MgCl_2_, 0.1% TritonX-100), fortified with 1X protease inhibitor (Halt protease inhibitor cocktail, Thermo, USA). Extracted protein samples were quantified by using Pierce™ BCA Protein Assay Kit (Thermo). A 30µg protein extracts were subjected to denaturing 10% SDS-polyacrylamide gels and transferred to polyvinylidene difluoride (PVDF) membrane preactivated with absolute methanol using Trans-Blot Turbo Transfer System (Bio-Rad). After blocking with 5% powdered skim milk in 1% phosphate buffer solution plus 0.1% Tween-20 (PBST), blots were probed with primary antibodies when included, anti-RbcL (PhytoAB; PHY5236A, a dilution of 1:2,000), PsbA (Agrisera; AS05084, a dilution of 1:5,000) and PsaC (Agrisera; AS10939, a dilution of 1:5,000) overnight at 4°C. After an hour incubation with secondary antibody a goat α-rabbit HRP conjugate (Invitrogen, G21234, a dilution of 1:20,000) at room temperature, antigen-antibody complexes were detected by enhanced chemiluminescence detection system (Super signal, Thermo scientific, USA). The Precision plus protein dual color standard (Bio-Rad) was used as reference molecular weight markers.

### Statistical analysis

Statistical analysis and plots were generated using MS excel, R and python. All experiments were performed with at least three biological replicates and technical replicates for same-day experiments are indicated. Data was shown as mean ± standard deviation (SD). Statistical analysis was performed using one-way ANOVA followed by Tukey’s multiple comparison test, or Student’s test and Mann-Whitney test with Benjamini-Hochberg correction, when appropriate. Differences were considered statistically significant at P < 0.05.

## Data availability

The mass spectrometry proteomics data have been deposited to the ProteomeXchange Consortium via the PRIDE (Perez-Riverol et al., 2019) partner repository with the dataset identifier PXD027430 and 10.6019/PXD027430. Proteins were searched against the uniprot protein database of UP000002717_Synehcococcus elongatus PC7942. Rubisco protein structure was downloaded from PDBe-KB and modified for figure display (Fig. 1A). The data that support the findings of this study are available from corresponding author on request.

## Acknowledgements

We would like to thank Dr. Curtis G. Wilkerson and Douglas Whitten for mass spectrometry analysis and critical feedbacks in experimental setup. We would like to thank Dr. Sigal Lechno-Yossef (Cheryl Lab) and Kaila Smith (Walker Lab) for technical assistance, and our lab pre-doctoral fellows Lisa Yun, Emmanuel Kokarakis and Rees Rillema for helpful comments on this manuscript. This work was supported by the Department of Energy, Basic Energy Sciences Division (Grant: DE-FG02-91ER20021).

